# The small and efficient language network of polyglots and hyper-polyglots

**DOI:** 10.1101/713057

**Authors:** Olessia Jouravlev, Zachary Mineroff, Idan A. Blank, Evelina Fedorenko

## Abstract

Acquiring a foreign language is challenging for many adults. Yet certain individuals choose to acquire sometimes dozens of languages, and often just for fun. Is there something special about the minds and brains of such polyglots? Using robust individual-level markers of language activity, measured with fMRI, we compared native language processing in polyglots versus matched controls. Polyglots (n=17, including 9 “hyper-polyglots” with proficiency in 10-55 languages) used fewer neural resources to process language: their activations were smaller in both magnitude and extent. This difference was spatially and functionally selective: the groups were similar in their activation of two other brain networks – the multiple demand network and the default mode network. We hypothesize that the activation reduction in the language network is experientially driven, such that the acquisition and use of multiple languages makes language processing generally more efficient. However, genetic and longitudinal studies will be critical to distinguish this hypothesis from the one whereby polyglots’ brains already differ at birth or early in development. This initial characterization of polyglots’ language network opens the door to future investigations of the cognitive and neural architecture of individuals who gain mastery of multiple languages, including changes in this architecture with linguistic experiences.

## Introduction

In the presence of linguistic input, a typically developing child effortlessly acquires a language, or multiple languages. Sometime during late childhood / early adolescence, after the so-called “critical period”, acquiring new languages becomes substantially more difficult (e.g., Lenneberg, 1967; Johnson & Newport, 1989; Pinker, 2009; Hyltenstam & Abrahamsson, 2003; Birdsong, 2005; DeKeyser & Larson-Hall, 2005; Hartshorne et al., 2018). Yet, many individuals learn new languages in their adult years, with some of them doing so of their own volition, suggesting they find the process enjoyable (e.g., Erard, 2012). *Polyglots* are a subset of these individuals, who obtain proficiency in multiple languages, sometimes dozens (“hyper-polyglots”; Hudson, 2016). Definitions of polyglotism vary in the literature, but most seem to reserve this term for individuals who have acquired at least some of the languages after the critical period (cf. multilinguals, who grow up in environments where multiple languages are spoken, like Belgium, Singapore, India, Morocco, or Mali) (e.g., Krzeminska, 2016; Hyltenstam, 2016).

Extraordinary cases of polyglotism have been reported (e.g., Baker & Prys Jones, 1997; Erard, 2012; Krashen & Kiss, 1996; Tyrkova-Williams, 1935), and modern-day polyglots continue to attract enormous attention, as evidenced by millions of views that videos of and about polyglots receive (e.g., Doner, 2014; Machova, 2018; THNKR, 2013), and by the high readership of popular articles about polyglots (e.g., Leland, 2012; Thurman, 2018). However, the cognitive, neural, and – to the extent innate predispositions exist – genetic bases of polyglotism remain poorly understood (e.g., Erard, 2012; Biedron & Pawlak, 2016; Hyltenstam, 2016). Are the minds and brains of polyglots different from those of monolingual individuals, or individuals who acquired multiple languages in early childhood? And to the extent that differences exist, are they restricted to language processing mechanisms or do they extend to other cognitive systems? We here begin to tackle these questions.

To the best of our knowledge (e.g., Hyltenstam, 2018), the only prior study that has asked this question was a post-mortem examination of the brain of Emile Krebs (E.K.), a German polyglot, who studied 120 languages and allegedly mastered over 60. Amunts et al. (2004) examined the microanatomy of Broca’s area (Brodmann areas 44 and 45) in E.K. compared to 11 male control brains and observed reliable cytoarchitectonic differences, with no differences observed in a control, visual, area (BA 18). However, two factors make the observed differences difficult to interpret. First, the features examined have not been linked to functional brain responses or behavior, which is not surprising given that this level of structural detail is not accessible to current imaging methodologies for living brains. And second, the postmortem nature of the study precludes matching the polyglot and control participants on cognitive abilities, like general IQ. The latter is especially important given that Broca’s area houses both language-selective and domain-general areas that have been linked to fluid intelligence (e.g., Fedorenko et al., 2012; Woolgar et al., 2018).

In the current study, we used functional MRI to probe the brains of 17 polyglots (languages spoken: 5-55), nine of whom qualified as “hyper-polyglots”, by Erard’s (2012) definition, having some knowledge of 10 or more languages. The polyglots were compared to pairwise-matched (on age, sex, handedness, and IQ) non-polyglots, as well as a larger control population (n=217). We examined activity during native language processing (English) in the fronto-temporal language network, which selectively supports high-level language comprehension, including both lexico-semantic and syntactic processing (e.g., Fedorenko et al., 2010, 2012, 2018). We also examined two control networks that support high-level cognition: the fronto-parietal domain-general multiple demand (MD) network (e.g., Duncan, 2010, 2013), which has been linked to executive control and fluid intelligence, and the fronto-parietal default mode network (e.g., Buckner et al., 2008), implicated in social cognition, recollection and prospection, and semantic processing.

We asked two research questions. First, we asked whether the language network differs between polyglots and controls, focusing on neural markers that we have previously established to be stable within individuals over time, like the strength and extent of activation, and lateralization (Mahowald & Fedorenko, 2016). A priori, one can make two opposite predictions. On the one hand, polyglots might exhibit stronger and more extensive language activations, perhaps reflecting richer/deeper processing of meaning and structure, in line with the general depth of processing idea (Nyberg, 2002), and with prior findings of larger anatomical structures (e.g., Maguire et al., 2003) or more extensive activations (e.g., Schneider et al., 2002; Maguire et al., 1997; Olesen et al., 2004; Russel et al., 2010) in experts in some domains. Alternatively, in line with prior work on activation reduction as a function of practice with a task (e.g., Poldrack et al., 1998; Fletcher et al., 1999; Kelly & Garavan, 2004; Bernardi et al., 2013), polyglots might exhibit weaker and less extensive activations, reflecting greater efficiency. Further, reduced lateralization of language processing has been reported in several language disorders (e.g., Sommer et al., 2001; Kleinhans et al., 2008; Bishop, 2013). If polyglotism is characterized by an aptitude for language learning, perhaps it would be associated with *increased* lateralization (e.g., Gotts et al., 2013; Mellet et al., 2014; cf. Novoa et al., 1988; Amunts et al., 2004).

And second, we asked whether the between-population differences are restricted to the language network or present in other networks that support high-level cognition. Bilingualism and multilingualism, at least in cases of acquisition within the critical period, have been associated with superior executive (e.g., Bialystok, 2009; Bialystok et al., 2005; cf. Paap & Greenberg, 2013; Duñabeitia et al., 2014), and mentalizing (Theory of Mind) abilities (e.g., Goetz, 2003; Kovacs, 2009). As a result, one might expect the differences to extend beyond the core fronto-temporal language network.

## Methods

### Participants

#### Polyglots

Polyglots were defined as individuals who (a) have some level of proficiency in at least five languages (their native language and four other languages), and (b) have advanced proficiency in at least one language other than their native language. Because this population has not been studied extensively in the past, these criteria are necessarily somewhat arbitrary. They are based on the fact that most individuals living in a predominantly monolingual society, like the US, typically study just one foreign language in school and/or college. So, having some proficiency in four foreign languages is sufficiently unusual. Participants assessed their own proficiency in listening, speaking, reading, and writing in each language they have some familiarity with, on a scale from 0=no knowledge to 5=native proficiency. A total score of 16 or higher for a language was used as an indicator of advanced proficiency. Seventeen polyglots were recruited from the Boston community (*M*_age_=30.5 years (*SD*=8.6); 9 males; 16 right-handed; all native speakers of English; *M*_*KBIT non-verbal*_=124 (*SD*=8)). The median number of languages spoken with some level of proficiency was 7 (*M*_languages_=13.9, range: 5-55 languages; see Table 1 for detailed linguistic background). Nine of the polyglots qualified as “hyper-polyglots”, having some knowledge of ten or more languages (Erard, 2012). The mean self-rated proficiency for L1 (native language) was 20.0 (*SD*=0), as expected, for L2 – 17.7 (*SD*=1.7, range: 16-20), for L3 – 15.84 (*SD*=2.52, range: 11-20), for L4 – 12.5 (*SD*=3.9, range: 6-20), and for L5 – 9.9 (*SD*=3.7, range: 4-16). Thus, in addition to having native-like proficiency in their L2, most of these individuals had quite high proficiency in their L3 and L4, and some in their L5. All polyglots were born in the US. Eleven polyglots were raised in monolingual households, while six grew up in bilingual families. Sixty percent of non-native languages spoken by polyglots were learned by them on their own using various self-learning tools (e.g., language learning software, textbooks, audio programs; see Hyltenstam (2018) for a discussion of “learner autonomy” in polyglots). Thirty one percent of non-native languages were acquired through language classes. The remaining nine percent were learned through immersion (3% in childhood through exposure to languages spoken by parents and family members; 6% in adulthood through travel to foreign countries).

**Table 1.**
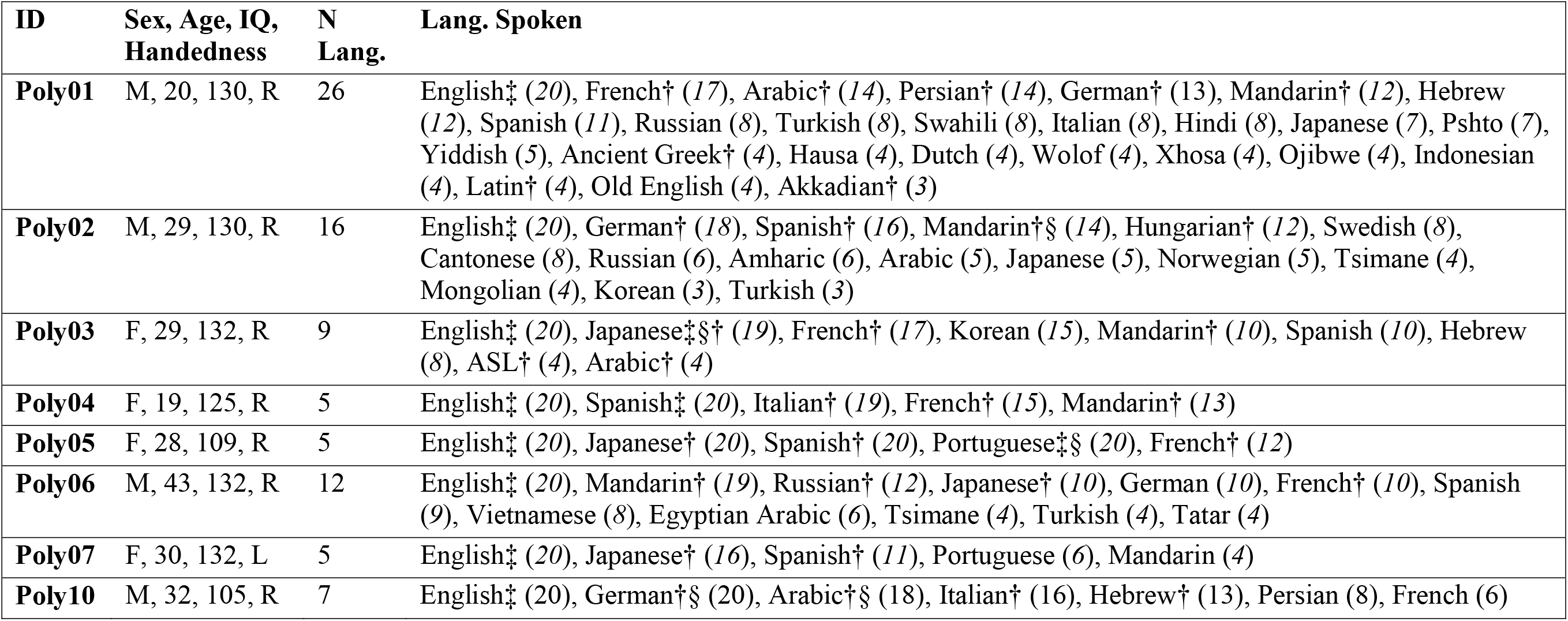
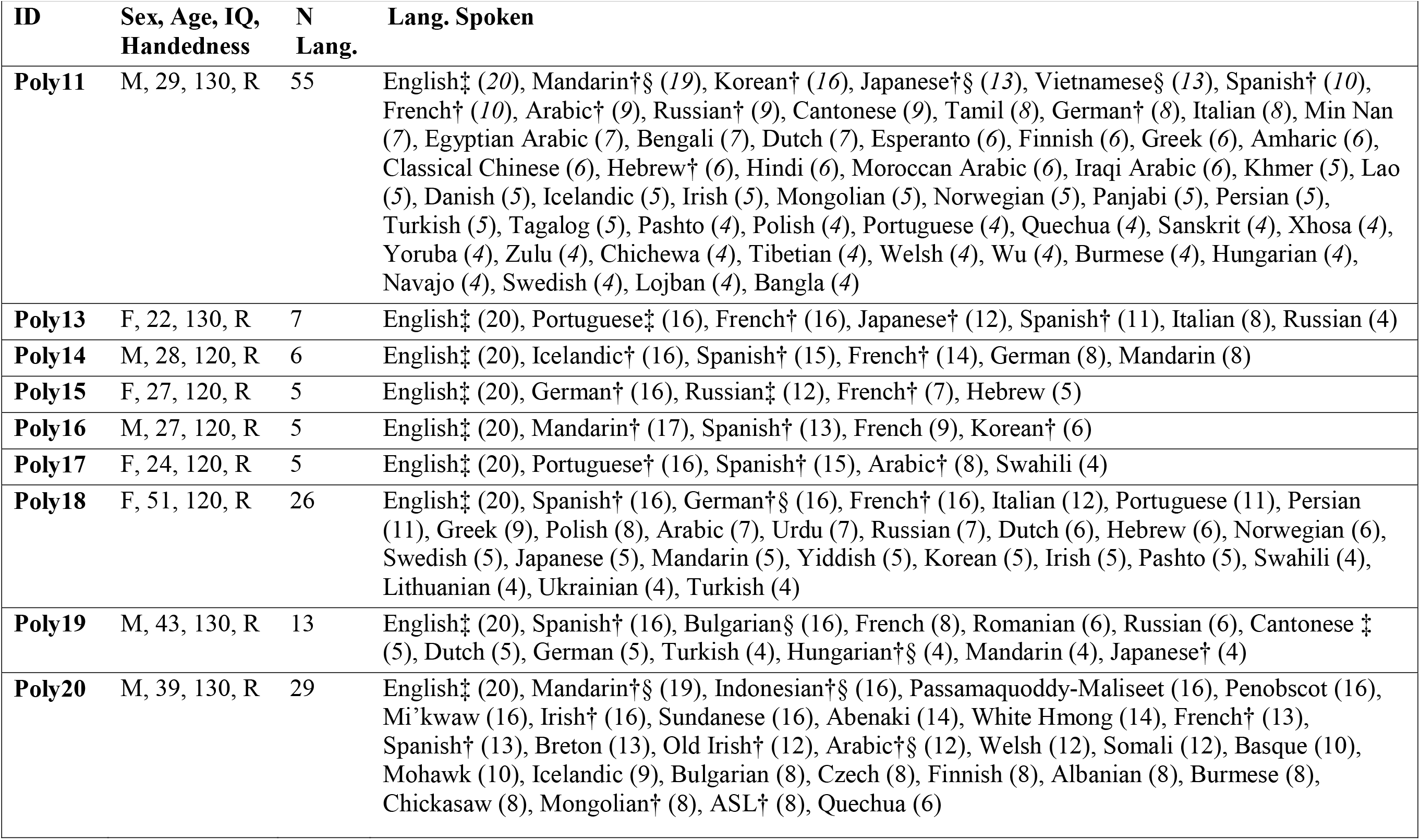
Demographic and linguistic background information for the sample of polyglots. Self-reported language proficiency scores are provided in brackets next to the names of languages (max. score of 20 corresponds to native proficiency). Symbols next to the names of languages indicate ways these languages were learned (‡-parents/immersion as a child; § - immersion as an adult; † - formal training, no symbol – self-training)

| ID | Sex, Age, IQ, Handedness | N Lang. | Lang. Spoken |
| --- | --- | --- | --- |
| **Poly01** | M, 20, 130, R | 26 | English‡ (20), French† (17), Arabic† (14), Persian† (14), German† (13), Mandarin† (12), Hebrew (12), Spanish (11), Russian (8), Turkish (8), Swahili (8), Italian (8), Hindi (8), Japanese (7), Pshto (7), Yiddish (5), Ancient Greek† (4), Hausa (4), Dutch (4), Wolof (4), Xhosa (4), Ojibwe (4), Indonesian (4), Latin† (4), Old English (4), Akkadian† (3) |
| **Poly02** | M, 29, 130, R | 16 | English‡ (20), German† (18), Spanish† (16), Mandarin†§ (14), Hungarian† (12), Swedish (8), Cantonese (8), Russian (6), Amharic (6), Arabic (5), Japanese (5), Norwegian (5), Tsimane (4), Mongolian (4), Korean (3), Turkish (3) |
| **Poly03** | F, 29, 132, R | 9 | English‡ (20), Japanese‡§† (19), French† (17), Korean (15), Mandarin† (10), Spanish (10), Hebrew (8), ASL† (4), Arabic† (4) |
| **Poly04** | F, 19, 125, R | 5 | English‡ (20), Spanish‡ (20), Italian† (19), French† (15), Mandarin† (13) |
| **Poly05** | F, 28, 109, R | 5 | English‡ (20), Japanese† (20), Spanish† (20), Portuguese‡§ (20), French† (12) |
| **Poly06** | M, 43, 132, R | 12 | English‡ (20), Mandarin† (19), Russian† (12), Japanese† (10), German (10), French† (10), Spanish (9), Vietnamese (8), Egyptian Arabic (6), Tsimane (4), Turkish (4), Tatar (4) |
| **Poly07** | F, 30, 132, L | 5 | English‡ (20), Japanese† (16), Spanish† (11), Portuguese (6), Mandarin (4) |
| **Poly10** | M, 32, 105, R | 7 | English‡ (20), German†§ (20), Arabic†§ (18), Italian† (16), Hebrew† (13), Persian (8), French (6) |

**Table 1 (continued).**
| **ID** | **Sex, Age, IQ, Handedness** | **N Lang.** | **Lang. Spoken** |
| --- | --- | --- | --- |
| **Poly11** | M, 29, 130, R | 55 | English‡ (20), Mandarin†§ (19), Korean† (16), Japanese†§ (13), Vietnamese§ (13), Spanish† (10), French† (10), Arabic† (9), Russian† (9), Cantonese (9), Tamil (8), German† (8), Italian (8), Min Nan (7), Egyptian Arabic (7), Bengali (7), Dutch (7), Esperanto (6), Finnish (6), Greek (6), Amharic (6), Classical Chinese (6), Hebrew† (6), Hindi (6), Moroccan Arabic (6), Iraqi Arabic (6), Khmer (5), Lao (5), Danish (5), Icelandic (5), Irish (5), Mongolian (5), Norwegian (5), Panjabi (5), Persian (5), Turkish (5), Tagalog (5), Pashto (4), Polish (4), Portuguese (4), Quechua (4), Sanskrit (4), Xhosa (4), Yoruba (4), Zulu (4), Chichewa (4), Tibetan (4), Welsh (4), Wu (4), Burmese (4), Hungarian (4), Navajo (4), Swedish (4), Lojban (4), Bangla (4) |
| **Poly13** | F, 22, 130, R | 7 | English‡ (20), Portuguese‡ (16), French† (16), Japanese† (12), Spanish† (11), Italian (8), Russian (4) |
| **Poly14** | M, 28, 120, R | 6 | English‡ (20), Icelandic† (16), Spanish† (15), French† (14), German (8), Mandarin (8) |
| **Poly15** | F, 27, 120, R | 5 | English‡ (20), German† (16), Russian‡ (12), French† (7), Hebrew (5) |
| **Poly16** | M, 27, 120, R | 5 | English‡ (20), Mandarin† (17), Spanish† (13), French (9), Korean† (6) |
| **Poly17** | F, 24, 120, R | 5 | English‡ (20), Portuguese† (16), Spanish† (15), Arabic† (8), Swahili (4) |
| **Poly18** | F, 51, 120, R | 26 | English‡ (20), Spanish† (16), German†§ (16), French† (16), Italian (12), Portuguese (11), Persian (11), Greek (9), Polish (8), Arabic (7), Urdu (7), Russian (7), Dutch (6), Hebrew (6), Norwegian (6), Swedish (5), Japanese (5), Mandarin (5), Yiddish (5), Korean (5), Irish (5), Pashto (5), Swahili (4), Lithuanian (4), Ukrainian (4), Turkish (4) |
| **Poly19** | M, 43, 130, R | 13 | English‡ (20), Spanish† (16), Bulgarian§ (16), French (8), Romanian (6), Russian (6), Cantonese ‡ (5), Dutch (5), German (5), Turkish (4), Hungarian†§ (4), Mandarin (4), Japanese† (4) |
| **Poly20** | M, 39, 130, R | 29 | English‡ (20), Mandarin†§ (19), Indonesian†§ (16), Passamaquoddy-Maliseet (16), Penobscot (16), Mi'kwaw (16), Irish† (16), Sundanese (16), Abenaki (14), White Hmong (14), French† (13), Spanish† (13), Breton (13), Old Irish† (12), Arabic†§ (12), Welsh (12), Somali (12), Basque (10), Mohawk (10), Icelandic (9), Bulgarian (8), Czech (8), Finnish (8), Albanian (8), Burmese (8), Chickasaw (8), Mongolian† (8), ASL† (8), Quechua (6) |

#### Matched monolingual controls

Polyglot participants were pairwise-matched with non-polyglots on age (*M* polyglot: 30.5 (*SD*=8.6) vs. *M* non-polyglot: 31.6 (*SD*=10.1); *t*(32)=0.33, *n.s.*), sex (9 males in each group), handedness (1 left-handed individual in each group), and nonverbal IQ, as measured by KBIT (Kaufman & Kaufman, 2004; *M* polyglot: 124 (*SD*=8) vs. *M* non-polyglot: 119.6 (*SD*=7.2); *t*(32)=1.74, *n.s.*). The mean number of languages with any level of proficiency for the non-polyglot controls was 1.4 (*SD*=0.5, range: 1-2). All non-polyglot controls with some knowledge of a second language identified as novice L2 speakers, and thus qualify as monolinguals.

#### A larger group of controls

To examine the key neural measures relative to a larger distribution from the population, we further included data from a relatively large (n=217) set of non-polyglot participants from the Fedorenko lab’s database each of whom had completed a language localizer experiment (Fedorenko et al., 2010) as part of different studies (*M*_*age*_=23.8 years (*SD*=6.1); 73 males; 205 right-handed; *M*_*KBIT*_=119.5 (*SD*=11.3); all native speakers of English; mean number of languages spoken with any level of proficiency=2.9 (*SD*=1.3, range: 1-9). In this dataset, individuals who spoke 5 or more languages did not have advanced proficiency in any of their non-native languages and thus do not qualify as polyglots. The 17 individuals that were pairwise-matched to the polyglots were excluded from this larger set.

All participants gave informed consent in accordance with the requirements of the Committee on the Use of Humans as Experimental Subjects at MIT. All participants were paid for their participation.

### Experimental design, materials and procedure

Each participant completed a language localizer task (Fedorenko et al., 2010) and a localizer for the Multiple Demand (MD) network (Duncan, 2010, 2013), which can also be used to define the regions of the Default Mode Network (e.g., Mineroff et al., 2018). Some participants completed one or two additional tasks for unrelated studies. The entire scanning session lasted approximately 2 hours.

#### Language localizer

The polyglots and the pairwise-matched non-polyglots passively read English sentences and lists of pronounceable nonwords in a blocked design. The Sentences>Nonwords contrast targets brain regions sensitive to high-level linguistic processing, including lexico-semantic and syntactic processes (Fedorenko et al., 2010, 2012, 2018) and has been shown to be robust to changes in materials, task, timing parameters, and other aspects of the procedure (Fedorenko et al., 2010; Fedorenko, 2014; Mahowald & Fedorenko, 2016; Scott, Gallée, & Fedorenko, 2017). Each trial started with 100 ms pre-trial fixation, followed by a 12-word-long sentence or a list of 12 nonwords presented on the screen one word/nonword at a time at the rate of 450 ms per word/nonword. Then, a line drawing of a hand pressing a button appeared for 400 ms, and participants were instructed to press a button whenever they saw this icon, and finally a blank screen was shown for 100 ms, for a total trial duration of 6 s. The simple button-pressing task was included to help participants stay awake and focused. Each block consisted of 3 trials and lasted 18 s. Each run consisted of 16 experimental blocks (8 per condition), and five fixation blocks (14 s each), for a total duration of 358 s (5 min 58 s). Each participant performed two runs. Condition order was counterbalanced across runs. One hundred sixty-eight of the 217 participants in the larger set performed the same version of the localizer task. The remaining 51 performed versions that differed slightly in the timing and other aspects of the procedure that have been shown to not affect the activations (e.g., Mahowald & Fedorenko, 2016). Information on the subsets of participants in the sample of 217 who completed different versions of the language localizer and details on procedure and timing for different versions of the language localizer is provided in Table 2.

**Table 2.** Information on which subsets of participants in the sample of 217 non-polyglot participants performed which version of the language localizer. Detailed information on procedure and timing details for the SNloc_ips179 is provided in the Methods section. Information on procedure and timing details for the other versions of the language localizer can be found in Mahowald & Fedorenko (2016), Table 2.

| Number of participants | Language localizer version | Conditions | Materials | Trials per block | Blocks per run / per condition per run |
| --- | --- | --- | --- | --- | --- |
| n = 168 | SNloc_ips179 | Sentences, Nonwords | 12-word-/nonword-long sequences | 3 | 16/8 |
| n = 29 | SNloc_ips189 | Sentence, Nonwords | 12-word-/nonword-long sequences | 3 | 16/8 |
| n = 20 | SWNloc_ips198 | Sentences, Wordlists, Nonwords | 12-word-/nonword-long sequences | 3 | 18/6 |
| n = 2 | SWJN_v1_ips252 | Sentences, Wordlists, Jaberwocky, Nonwords | 12-word-/nonword-long sequences | 5 | 16/4 |
| n = 1 | SWJN_v2_ips232 | Sentences, Wordlists, Jaberwocky, Nonwords | 8-word-/nonword-long sequences | 5 | 16/4 |
| n = 1 | SNloc_ips232 | Sentences, Nonwords | 8-word-/nonword-long sequences | 5 | 16/8 |

#### Multiple Demand and Default Mode network localizer

Participants performed a spatial working memory task, where they had to keep track of four (easy condition) or eight (hard condition) sequentially presented locations in a 3 x 4 grid (Blank et al., 2014). In both conditions, subjects performed a two-alternative forced-choice task at the end of each trial to indicate the set of locations that they just saw. The Hard>Easy contrast targets the brain regions of the Multiple Demand (MD) network, a bilateral fronto-parietal network that supports executive functions, is broadly engaged across domains, and is modulated by effort (Duncan, 2010, 2013; Fedorenko et al., 2013). The reverse, Easy>Hard, contrast can be used to identify the brain regions of the Default Mode Network (DMN), a bilateral network that has been linked to internally-directed cognition (Buckner et al., 2008), because a well-established functional signature of this network is deactivation to demanding tasks, with greater deactivation to more demanding conditions. Indeed, DMN regions defined with the Easy>Hard contrast from the spatial working memory task show exactly this profile (Mineroff et al., 2018), and contrasts between fixation and the Easy or Hard condition yield similar areas.

Each trial lasted 8 s (see Fedorenko et al., 2011, for the timing details). Each block consisted of 4 trials and lasted 32 s. Each run consisted of 12 experimental blocks (6 per condition), and 4 fixation blocks (16 s in duration each), for a total duration of 448 s (7 min 28 s). Sixteen polyglots and 16 matched controls performed two runs of the task; the remaining 1 participant in each group performed one run of the task. In the large set of non-polyglots, 165 participants completed two runs of the task; the remaining 52 performed one run of the task. Condition order was counterbalanced across runs when participants performed two runs. Only the data of participants who completed two runs of the task were used to examine group differences in the MD and DMN network activity (because across-runs cross-validation is necessary to maintain independence between the data used to define the fROIs and to characterize their responses).

### fMRI data acquisition, preprocessing, and modeling

Structural and functional data were collected on the whole-body 3 Tesla Siemens Trio scanner with a 32-channel head coil at the Athinoula A. Martinos Imaging Center at the McGovern Institute for Brain Research at MIT. T1-weighted structural images were collected in 179 sagittal slices with 1 mm isotropic voxels (TR=2530ms, TE=3.48ms). Functional, blood oxygenation level dependent (BOLD) data were acquired using an EPI sequence (with a 90° flip angle and using GRAPPA with an acceleration factor of 2), with the following acquisition parameters: thirty-one 4mm thick near-axial slices, acquired in an interleaved order with a 10% distance factor; 2.1mm × 2.1mm in-plane resolution; field of view of 200mm in the phase encoding anterior to posterior (A>P) direction; matrix size of 96 × 96 voxels; TR of 2000ms; and TE of 30ms. Prospective acquisition correction (Thesen et al., 2000) was used to adjust the positions of the gradients based on the participant’s motion one TR back. The first 10s of each run were excluded to allow for steady-state magnetization.

fMRI data were preprocessed and analyzed (for basic data modeling) in SPM5 and custom MATLAB scripts. (Note that preprocessing and basic modeling have not changed much in the later versions of SPM, as confirmed by direct comparisons on several datasets performed in our lab; we chose to use the older version here because the data for some of the control participants were collected and analyzed many years ago, and we wanted to have all the data analyzed through the same pipeline, for better comparability.) Each subject’s data were motion corrected and then normalized into a common brain space (the Montreal Neurological Institute (MNI) template) and resampled into 2mm isotropic voxels. The data were then smoothed with a 4mm Gaussian filter and high-pass filtered (at 200s). For each localizer task, a standard mass univariate analysis was performed whereby a general linear model estimated the effect size of each condition in each experimental run. These effects were each modeled with a boxcar function (representing entire blocks) convolved with the canonical hemodynamic response function. The model also included first-order temporal derivatives of these effects, as well as nuisance regressors representing entire experimental runs and offline-estimated motion parameters.

### Language, Multiple Demand (MD), and Default Mode Network (DMN) fROI definition and estimation of the neural features of interest

For each participant, functional regions of interest (fROIs) were defined using the Group-constrained Subject-Specific (GSS) approach (Fedorenko et al., 2010), whereby a set of parcels or “search spaces” (i.e., brain areas within which most individuals in prior studies showed activity for the localizer contrast; Figure 1) is combined with each individual participant’s activation map for the same contrast.

**Figure 1.**
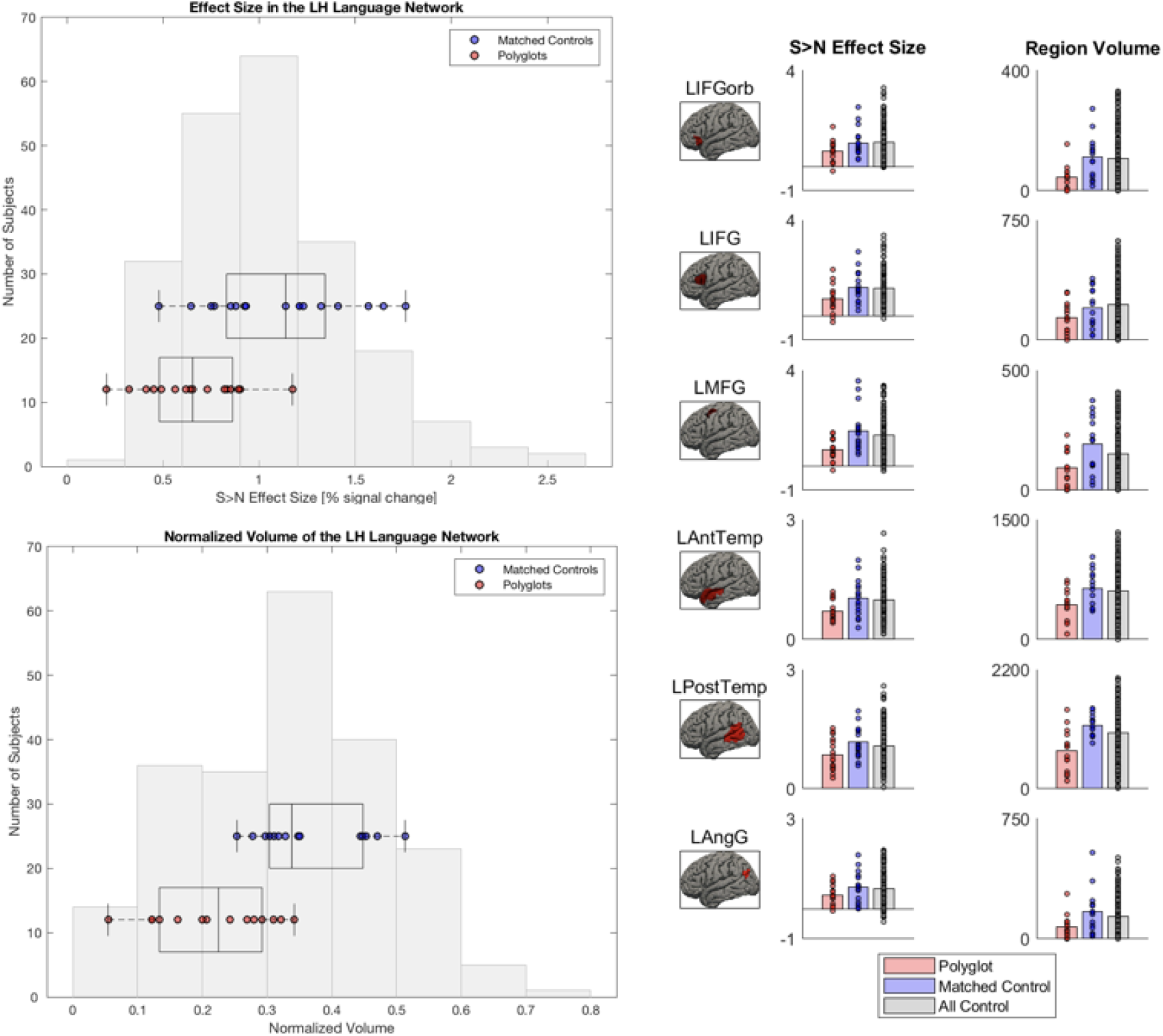
Functional properties of the LH language network in polyglots vs. non-polyglots. Left: The *Sentences>Nonwords* effect sizes and region volumes are shown as box-and-whisker plots for polyglots and matched non-polyglots (red and blue dots, respectively) and as a histogram for a larger sample of non-polyglots. Right: The effect sizes and region volumes for the three groups (polyglots, matched non-polyglots, larger set of non-polyglots) are shown as bar plots for the six language regions separately. Group differences in effect sizes and region volumes were present in the LH language network.

To define the language fROIs, we used six parcels derived from a group-level representation of data for the Sentences>Nonwords contrast in 220 participants. These parcels included three regions in the left frontal cortex: two located in the inferior frontal gyrus (LIFG and LIFGorb), and one located in the middle frontal gyrus (LMFG); and three regions in the left temporal and parietal cortices spanning the entire extent of the lateral temporal lobe and extending into the angular gyrus (LAntTemp, LPostTemp, and LAngG). Additionally, we examined activations in the right hemisphere homologs of the language regions. To define the fROIs in the right hemisphere, the left hemisphere parcels were mirror-projected onto the RH to create six homologous masks. By design, the parcels cover large swaths of cortex in order to be able to accommodate inter-individual variability. Hence the mirrored versions are likely to encompass RH language regions despite possible hemispheric asymmetries in the precise locations of activations (Mahowald & Fedorenko, 2016).

To define the MD fROIs, we used eighteen anatomical parcels in the frontal and parietal cortices of the two hemispheres (Tzourio-Mazoyer et al., 2002). These regions included bilateral opercular IFG (L/R IFGop), MFG (L/R MFG), orbital MFG (L/R MFGorb), insular cortex (L/R Insula), precentral gyrus (L/R PrecG), supplementary and presupplementary motor areas (L/R SMA), inferior parietal cortex (L/R ParInf), superior parietal cortex (L/R ParSup), and anterior cingulate cortex (L/R ACC). (These anatomical parcels are highly overlapping with a set of functional parcels derived from a group-level representation of data for the Hard>Easy spatial working memory contrast in 197 participants. We chose to use the anatomical parcels here for consistency with prior studies (Fedorenko et al., 2013; Blank et al., 2014).)

To define the DMN fROIs, we used eight anatomical parcels in the frontal and parietal cortices of the two hemispheres. These regions included posterior cingulate (L/R PostCing), frontal medial orbital cortex (L/R FrontMedOrb), frontal medial superior cortex (L/R FrontMedSup), and the precuneus (L/R Precuneus). In addition, we included two parcels – in the left and right temporo-parietal junction (L/R TPJ) – derived from a group-level representation of data for the False Belief>False Photograph contrast in 462 participants (Dufour et al., 2013). (This set of 10 parcels are highly overlapping with a set of functional parcels derived from a group-level representation of data for the Easy>Hard spatial working memory contrast in 197 participants.)

We examined three features of the activations for our four networks of interest (left hemisphere (LH) language, right hemisphere (RH) language, MD, and DMN): i) effect sizes, ii) extent of activation (region volumes), and iii) lateralization based on the extent of activation. All three measures have been shown to be reliable within individuals over time (e.g., Mahowald & Fedorenko, 2016; Assem et al., 2018). The first two measures have further been shown to be strongly correlated (e.g., Mahowald & Fedorenko, 2016); as a result, whatever differences emerge with respect to effect sizes are expected to also manifest for the extent of activation measures.

To compute effect sizes, individual fROIs were defined by selecting – within each parcel – the top 10% of most localizer-responsive voxels based on the *t* values for the relevant contrast (Sentences>Nonwords for the language network localizer, Hard>Easy spatial working memory for the MD network localizer, and Easy>Hard spatial working memory for the Default Mode Network localizer). To maintain independence between the data used to define the fROIs vs. to characterize their responses (Kriegeskorte et al., 2009), we used an across-run cross-validation procedure, where i) the first run was used to define the fROIs, and the second run to estimate the responses (in percent BOLD signal change); ii) the second run was used to define the fROIs, and the first run to estimate the responses; and finally, iii) the estimates were averaged across the two left-out runs to derive a single value per participant per fROI.

To compute region volumes, we counted the number of voxels that showed a significant effect (at the p<0.001 whole-brain uncorrected threshold) for the relevant localizer contrast within each parcel.

Finally, to estimate the degree of lateralization (for the language network only), we subtracted the number of activated voxels for the Sentences>Nonwords contrast (at the p<0.001 whole-brain uncorrected threshold) across all the right hemisphere language parcels from the number of Sentences>Nonwords voxels in all the left hemisphere parcels and divided the resulting value by the total number of Sentences>Nonwords voxels across hemispheres. The resulting lateralization values range from 1 (exclusively left-hemisphere activations) to −1 (exclusively right-hemisphere activations).

### Statistical Analyses

To test whether high-level language-processing regions differ in their functional properties between polyglots and non-polyglots, we used general linear models (GLMs) and Bayesian linear regressions with Group (Polyglots vs. Non-polyglots) as a predictor of the Sentences>Nonwords effect sizes (in the LH and RH separately), region volumes (in the LH and RH separately), and lateralization. Bayes Factor (BF10) statistics were calculated using the JASP software package (JASP Team, 2019). We did not correct the results for the use of three neural measures. First, as noted above, effect sizes and region volumes are strongly correlated (e.g., Mahowald & Fedorenko, 2016); as a result, we treated the region volume analyses as complementary to the effect size analyses, expecting them to mirror each other (which they did, at least at the network level). And second, the lateralization measure, albeit largely independent from the effect size / region volume measures, was used to evaluate a distinct hypothesis.

For each of the three measures, the GLM and Bayesian liner regression analyses described above were conducted across the language network. For the effect size and region volume measures, we further examined each of the 6 fROIs separately (correcting for the number of fROIs within each network), to test for potential differences among the regions. To examine the effects across the network, effect sizes were averaged across the regions within each network. Region volume measures were summed across the regions within each network and normalized by dividing the number of activated voxels by the total number of voxels in the network (i.e., 6,794 voxels total for each of the LH and RH language networks).

To circumvent the issue of a relatively small sample of polyglots, we also assessed the probability that the polyglots (or the matched non-polyglots, for comparison) were drawn from the same distribution as a relatively large population (n=217) of non-polyglots, for each neural measure. This was done via permutation tests, by randomly sampling (10,000 times) 17 data points from the large set of non-polyglots and comparing the observations from the polyglots (or the matched controls) to the distribution of these random samples. These analyses complement the critical analyses performed with the carefully pairwise matched controls.

Next, to test whether non-language brain networks differ in their functional properties between polyglots and non-polyglots, we used general linear models (GLMs) and Bayesian linear regressions with Group (Polyglots vs. Non-polyglots) as a predictor of (a) the Hard>Easy effect sizes and region volumes for the bilateral MD network, which supports executive functions (Duncan, 2010), and (b) the Easy>Hard effect sizes and region volumes for the bilateral DMN network, which supports internally-directed cognition (Buckner et al., 2008). All the analyses were parallel to those carried out on the LH and RH language networks above. For region volume normalization, the following values were used: 41,012 voxels total for the MD network, and 15,070 voxels total for the DMN network.

To test whether the patterns of polyglot vs. non-polyglot differences differed between the LH language network and the other networks examined, the key measures (effect sizes and region volumes for each relevant contrast) served as dependent variables in three linear mixed-effects models (performed using the lme4 package in R; Bates et al., 2014) which included a fixed effect for a Network (LH Language vs. Control, where Control was the RH Language network, the Multiple Demand network, or the Default Mode network) x Group (Polyglots vs. Non-polyglots) interaction, and random intercepts for participants. Significance values were obtained using the likelihood ratio tests. To compare the polyglots to the larger sample of non-polyglots, the same permutation tests were used as those described above.

## Results

### 1. The LH language network is smaller and less active in polyglots

The polyglots’ LH language network showed lower activation and was smaller in its extent compared to both the matched controls (*t*(32)=3.67, *p*<0.001, *d* = 1.36, *BF10*s=70; *t*(32)=4.04, *p*<0.001, *d*=1.57, *BF10*s=294; Figure 1; see Figure 2 for sample individual language activation maps in polyglots vs. controls) and the larger sample of non-polyglots (*ps*<0.001; Figure 1). The observed group differences were also reliable in most individual fROIs (Figure 1, right panel): the polyglots showed weaker responses than the controls in the LAntTemp, LPostTemp, LIFG, and LMFG fROIs (*ts*(32)>2.34, *ps*<0.03, *ds*>.86, FDR-corrected for the number of fROIs here and below; although note that in the Bayesian analyses, moderate or strong evidence for group differences in activation of the LH language network was found only for LAntTemp, LPostTemp, and LMFG fROIs, *BF10*s>3.26), and smaller region volumes in the LAntTemp, LPostTemp, LMFG, and LIFGorb fROIs (*ts*(32)>2.62, *ps*<0.02, *ds*>.88, *BF10*s>3.62). Further, the polyglots (but not the matched non-polyglots) showed reliably weaker responses and smaller regions relative to the larger sample of non-polyglots (*n*=217) for all language fROIs (*ps*<0.03).

**Figure 2.**
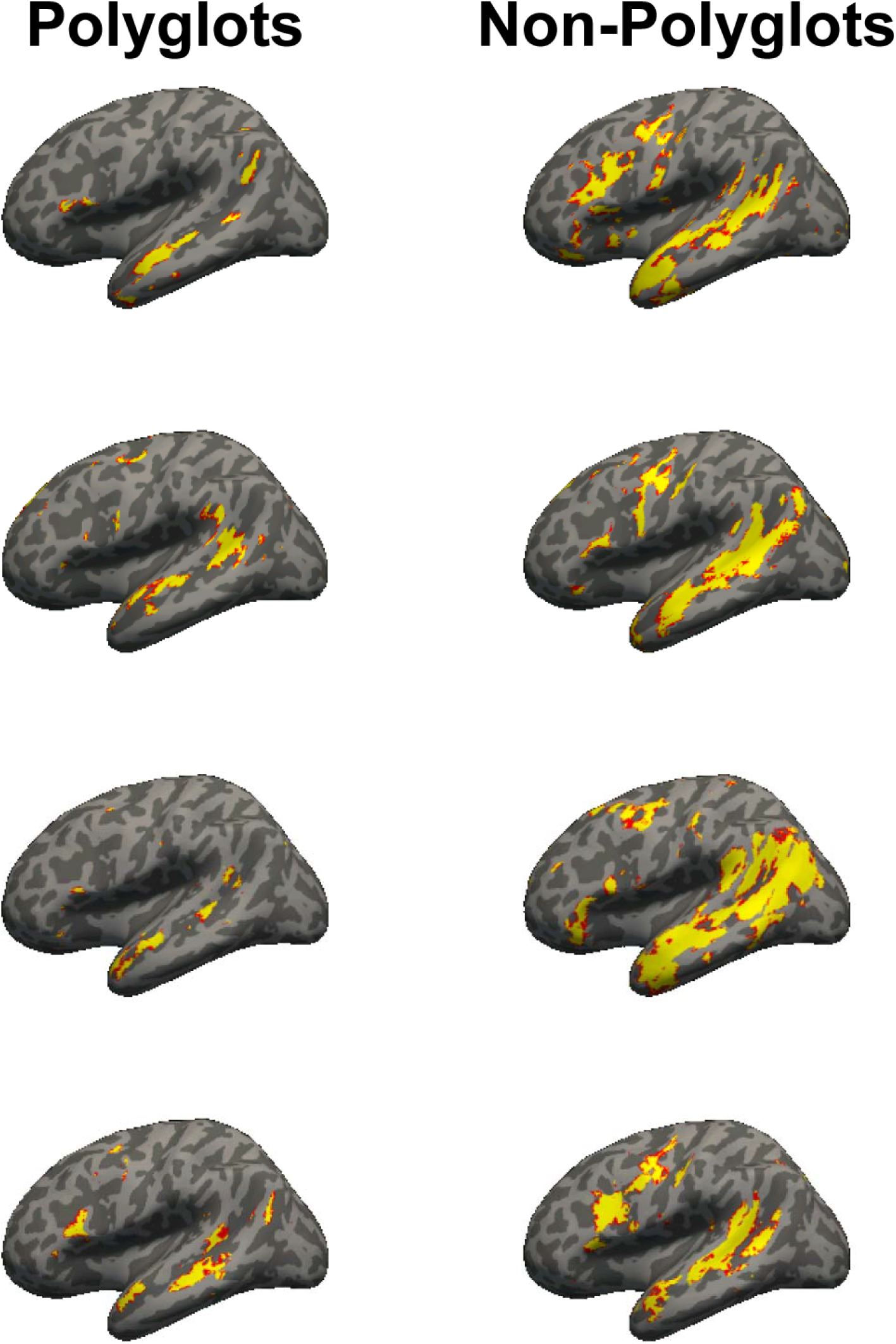
Whole-brain maps of language activity (at the threshold of p<0.001, uncorrected) in five polyglot: matched-control pairs. Polyglots exhibited weaker and less extensive activity.

### 2. Reduced neural activity in polyglots is restricted to the LH language network

There was no evidence that the strength or extent of activation in the RH language network differed between the polyglots and the matched controls (*ts*(32)<1, *p*s>0.52,*d*s<.22, *BF10*s<0.38; Figure 3) or the larger sample of non-polyglots (*ps*>0.26). Similarly, we found no evidence of group differences in two control domain-general networks (Figure 3): MD network (polyglots vs. matched controls: *ts*(30)<0.20, *ps*>0.80, *d*s<.08, *BF10*s<0.34; polyglots vs. a larger sample: *ps*<0.13) and DMN (polyglots vs. matched controls: *ts*(30)<0.69, *ps*>.49, *d*s<.26, *BF10s*<0.40; polyglots vs. a larger sample: *ps*<0.25). Further, the effects observed in the LH language network differed reliably from those observed in each of the other three networks (RH language, MD, DMN), as evidenced by reliable Network x Group interactions (χ^*2*^*s* (1)>4.11, *p*s<0.04).

**Figure 3.**
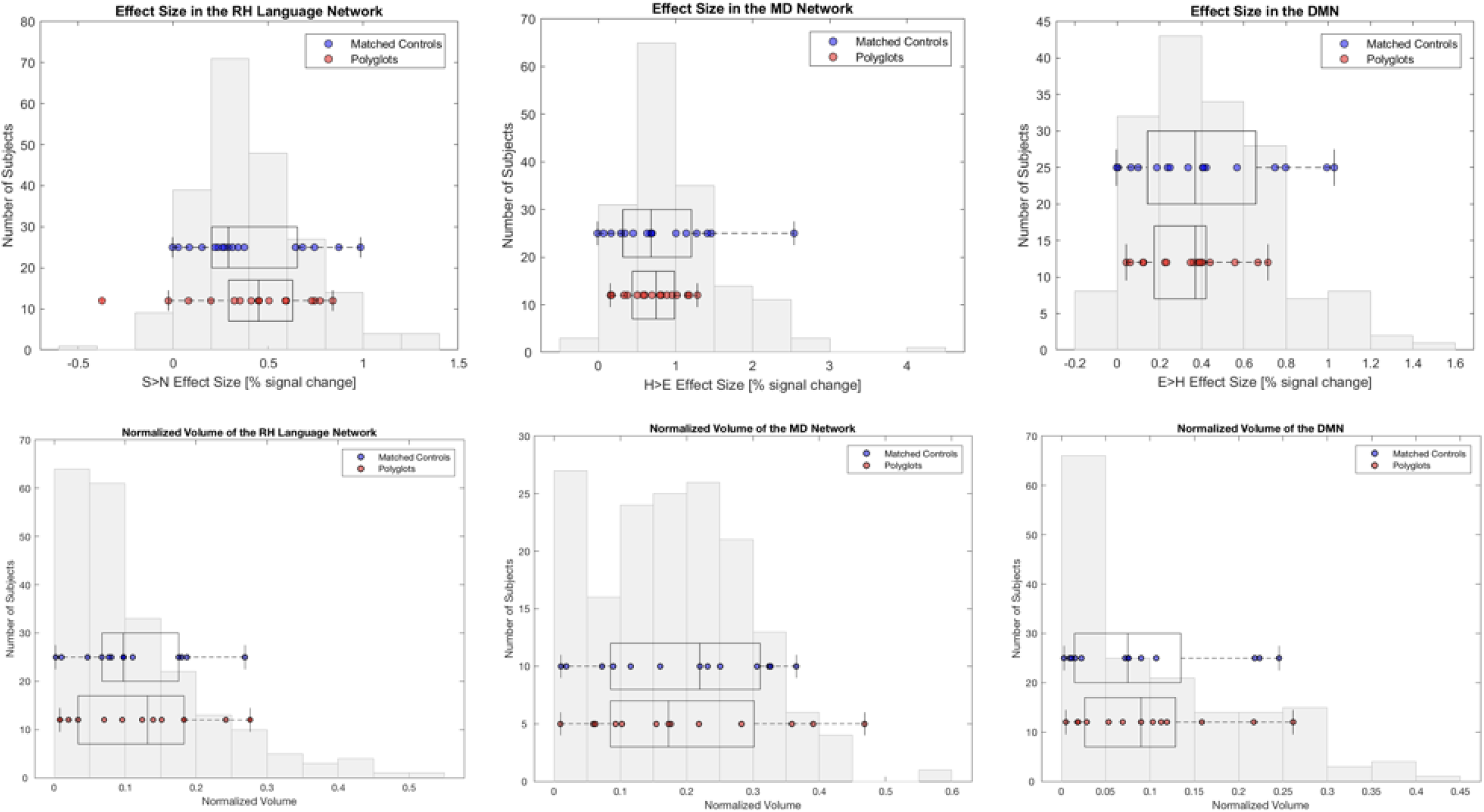
Functional properties of the RH Language, MD and DMN regions in polyglots vs. non-polyglots. The *Sentences>Nonwords* (Language), *Hard>Easy* (MD) and *Easy>Hard* (DMN) effect sizes and region volumes are shown as box-and-whisker plots for polyglots and matched non-polyglots (red and blue dots, respectively) and as a histogram for a larger sample of non-polyglots. No group differences were found in the RH Language network on in any of the control networks.

In the presence of similar strength and extent of activation in the RH language network between the polyglots and controls, the weaker LH language activations led to a significant group difference in the degree of language lateralization, with the polyglots exhibiting less lateralized responses (*t*(32)=2.51, *p*=0.02, *d*=.85, *BF10*=3.34). This result was corroborated by the permutation test that found less lateralized responses in the polyglots compared to the larger sample of non-polyglots (*p*=0.02).

## Discussion

Much past research has focused on developmental and acquired impairments that affect the acquisition and/or processing of language (e.g., Bloom & Lahey, 1978; Goodglass, 1993; Gorno-Tempini et al., 2011; Schwartz, 2017). Understanding how a cognitive system may break is a powerful approach that has yielded core insights into the architecture of the human mind. However, a complementary, and potentially similarly powerful, approach is to probe the minds and brains of individuals with special aptitude for a particular cognitive domain (e.g., Obler & Fein, 1988; Padgett & Seaberg, 2014; Russell et al., 2009; Winner, 1997). Linguistic aptitude / expertise can manifest in many ways, from an exceptionally large vocabulary (e.g., avid readers), to fast and eloquent speech (e.g., orators), to the ability to quickly come up with rhymes (e.g., rappers) or find the precise word or phrase to express an idea (e.g., journalists or novelists), to the extraordinary spelling ability (e.g., spelling bee champions). Another form of linguistic aptitude lies in the ability to learn multiple foreign languages after the critical period. Whether and how the minds and brains of linguistic experts, including polyglots, differ from those of typical language users remains poorly understood, yet might critically inform our understanding of language learning and processing.

This work is the first to characterize the functional properties of the language network in the brains of polyglots – individuals capable of communicating in five or more languages. To illuminate the neural architecture of the language system of polyglots, we conducted a cross-sectional fMRI study (see Poldrack, 2000, on the benefits and limitations of this approach) where we compared neural responses in the language network – and two control networks – of 17 polyglots (range of languages spoken: 5-55) with those of 17 carefully matched non-polyglot controls as well as a larger set of non-polyglots (n=217). Four results emerged clearly, as elaborated below.

**First**, the polyglots appeared to have a smaller language network. Compared to non-polyglot controls, polyglots recruited less extensive cortical areas within the fronto-temporal language network of the left hemisphere (reflected in smaller region volumes) and activated these areas to a lesser degree (reflected in smaller response magnitudes). These expertise-related differences in the properties of the language network are in line with prior reports of anatomical and functional changes in the brain in response to knowledge and skill acquisition (e.g., Bavelier et al., 2012; Calvo-Merino et al., 2004; Gauthier et al., 1999; Landau & d’Esposito, 2006; Maguire et al., 2003; McCandliss et al., 2003; Protzner et al., 2016; Schneider et al., 2002).

Neural differences between any group of experts and controls are notoriously difficult to interpret, however, because cross-sectional designs fail to determine whether the observed differences *result* from the extensive training or whether instead they initially *spur* some individuals to seek training in the relevant domain (e.g., Yarrow et al., 2009; Zatorre et al., 2012). Thus, reduced language activity in polyglots might reflect extensive linguistic experience: language representation and processing may become more efficient as a result of acquiring multiple languages. This would parallel activation reduction in other domains, like motor learning (e.g., Bernardi et al., 2013; Fletcher et al., 1999; Kelly & Garavan, 2005; Poldrack et al., 1998). Kelly and Garavan (2005) refer to this experientially-induced reduction of neural activity as a “processing efficiency change”. However, it is also possible that individuals who eventually become polyglots represent and process language more efficiently from the start, even as they acquire their first (native) language. Without establishing a genetic basis for polyglotism (Graham & Fisher, 2013), combined with longitudinal investigations of individuals as they acquire new languages (Osterhout et al., 2006), we cannot conclusively determine the causal direction of the observed group difference.

**Second**, the difference between the polyglots and controls was restricted to the language network in the dominant (left) hemisphere. Activations in the right-hemisphere homologs of the language regions were similar between the two groups. The role of the right hemisphere language network in linguistic/cognitive processing is widely debated (e.g., Gazzaniga & Hillyard, 1971; Jung-Beeman, 2005; Lindell, 2006; Vigneau et al., 2011). Our results do not inform this debate directly. However, the fact that the observed group differences were restricted to the left hemisphere adds to the evidence that the LH and RH language regions constitute complementary but distinct networks (e.g., Gotts et al., 2013; Chai et al., 2016) that can be differentially affected by linguistic experience, or possess distinct early, possibly genetically-driven, biases.

**Third**, the polyglots exhibited a reduced degree of left-lateralization for language compared to non-polyglots. Interestingly, reduced lateralization of linguistic function has been previously reported in numerous developmental disorders, including those characterized by language impairments, such as autism, specific language impairment, dyslexia, and schizophrenia (e.g., De Guilbert et al., 2011; Herbert et al., 2002; Oertel-Knochel & Linden, 2011; Wehner, Ahlfors, & Mody, 2007). Reduced language lateralization has been argued to index linguistic deficits or lower linguistic ability in neurotypical individuals (e.g., Bishop, 2013; Mellet et al., 2014). Our observation of reduced language lateralization in individuals with (at least one kind of) exceptional linguistic abilities appears to contradict this idea. However, the underlying causes of lateralization reduction in these different populations are distinct. In linguistic disorders, reduced lateralization is due to the greater engagement of the right hemisphere (e.g., Anderson et al., 2010; Jouravlev et al., submitted; Kleinhans et al., 2008; Takeuchi et al., 2004; Tesink et al., 2009), whereas in the polyglots, it results from reduced activity in the left hemisphere. Thus, reduced lateralization can apparently characterize both ends of the linguistic abilities spectrum: linguistic impairments (in the presence of increased RH activity) and linguistic aptitude/expertise (in the presence of reduced LH activity).

**Finally**, we found no evidence that polyglotism has a widespread effect across the brain. The strength and extent of activation were similar between the polyglots and controls in two domain-general brain networks linked to high-level cognition, including some aspects of language / communication – the Multiple Demand network, which supports executive functions (Duncan, 2010), and the Default Mode Network, which supports internally-directed cognition (Buckner et al., 2008). This result argues against ubiquitous between-group differences in information processing, and is in line with prior work that has suggested that the language network is functionally distinct from other high-level large-scale networks (e.g., Blank et al., 2014; Fedorenko et al., 2011; Fedorenko & Varley, 2016; Mineroff et al., 2018; Monti et al., 2012).

To conclude, compared to matched non-polyglots, as well as a larger population of control participants, the polyglots have smaller language regions that respond less strongly during native language processing. This difference is restricted to the left-hemisphere language network and may reflect more efficient processing in polyglots (Fletcher et al., 1999; Kelly & Garavan, 2004; Poldrack et al., 1998). However, the nature of this difference – innate/early-emerging vs. driven by the experience of acquiring multiple languages – requires further investigation, including longitudinal studies and studies that probe the genetic basis of polyglotism.

## Acknowledgements

We thank i) EvLab members for help with fMRI data collection and helpful comments, ii) Ted Gibson, Nancy Kanwisher, Simon Fisher, and Michael Erard for helpful discussions and/or comments on the manuscript, iii) Evgeniia Diachek for help with creating the figures, and iv) all our polyglots and other participants. The authors would also like to acknowledge the Athinoula A. Martinos Imaging Center at the McGovern Institute for Brain Research at MIT, and the support team (Steven Shannon and Atsushi Takahashi). E.F. was supported by NIH awards R00-HD057522, R01-DC016607, and R01-DC016950.

